# Environmental stress during larval development induces head methylome profile shifts in the migratory painted lady butterfly (*Vanessa cardui*)

**DOI:** 10.1101/2022.11.10.515964

**Authors:** Jesper Boman, Yishu Zhu, Lars Höök, Roger Vila, Gerard Talavera, Niclas Backström

## Abstract

Seasonal environmental fluctuations provide formidable challenges for living organisms, especially small ectotherms such as butterflies. A common strategy to cope with harsh environments is to enter diapause, but some species avoid the unsuitable conditions by migrating. Despite a growing understanding of migration in the life cycles of some butterfly species, it is unknown how individuals register and store environmental cues to determine whether and where to migrate. Here, we explored how competition and host plant availability during larval development affect patterns of DNA methylation in the migratory painted lady (*Vanessa cardui*) butterfly. We identify a set of potentially functional methylome shifts associated with differences in the environment, indicating that DNA methylation is involved in the response to different conditions during larval development. By analysing the transcriptome for the same samples used for methylation profiling, we also uncovered a non-linear relationship between gene body methylation and gene expression. Our results provide a starting point for understanding the interplay between DNA methylation and gene expression in butterflies in general and how differences in environmental conditions during development can trigger unique epigenetic marks that might be important for behavioural decisions in the adult stage.

## 1. Background

Organisms are forced to adapt to seasonal variation in both abiotic and biotic factors. Butterflies and moths (lepidopterans) have adopted strategies to cope with unfavourable conditions by either entering a state of dormancy and metabolic inactivity (diapause) or by migrating to regions with more favourable conditions, or a combination thereof [1,2]. While the mechanistic underpinnings of diapause in lepidopterans have been investigated in some detail [3–9], research on the molecular machinery underlying migratory behaviour is still in its infancy, except in the monarch butterfly (*Danaus plexippus*), for which some progress has been made on e.g. the neurogenetics of navigation [reviewed in 10]. During development, migrating insects face a trade-off between reproduction and migration [11]. For females, such a resource allocation trade-off predominantly involves temporal arrest of mating organs and egg development until a migratory flight has been completed, which is commonly referred to as the “oogenesis-flight” syndrome [11].

The painted lady (*Vanessa cardui*) is an emerging model system for investigating migratory behaviour in insects [12–20]. Besides being the most widespread of all butterflies – the species distribution is almost cosmopolitan – the painted lady is also the most long-distance migrant of all butterflies [19,21]. Painted ladies undertake an annual migratory cycle that can span ~ 6-10 generations. The most well studied circuit runs from tropical Africa to northern Europe and back [16,19,21]. Since the migratory movement of the painted lady is multi-generational, the direction and duration of migratory flight cannot be “learned” by conspecifics. Instead, individuals must use environmental cues to determine whether and where to migrate, but it is not known at what life stage the decisions are made. In captivity, female painted ladies mate on average 5.8 days after eclosion from pupae [12]. An important environmental cue is likely host plant abundance [22,23], which hypothetically could be sensed already in the early larval stages. Both the abundance of food and the density of competing conspecifics could inform the developing larvae about the expected future availability of host plants. For this signal to influence the onset of migratory behaviour of the adult butterfly after eclosion from pupa, the information needs to be stored, processed, and transmitted during metamorphosis.

One molecular mechanism that fulfils the requirements is epigenetic modifications, especially 5’-CpG-3’ DNA methylation (hereafter ‘methylation’), which can be transmitted to daughter strands during DNA replication with high fidelity [24]. In plants and vertebrates, methylation of promoter sequences has been linked to transcriptional repression [25–28], while the functional role of gene body methylation is still under debate [29–31]. In arthropods, patterns of methylation are less well characterized and potential functional relationships with transcription have been investigated to some extent but are inconclusive [32–40]. Recent initiatives suggest that the genome-wide levels of methylation can vary from moderate (~ 20%) in some arthropod species to almost undetectable (~ 0%) in others [34,35,41]. An analysis of silk gland tissue of silk moth (*Bombyx mori*) larvae revealed a low genome-wide methylation level (0.11%) and a positive relationship between gene expression and gene body methylation [32]. Another investigation of potential relationships between methylation and expression in adult cotton bollworm moths (*Helicoverpa armigera*) with different flight performance phenotypes pinpointed some differentially methylated CpG sites [38]. However, the study covered only 16 genes and the data were not sufficient to establish if differentially methylated sites were enriched in differentially expressed genes.

Here we investigate the DNA methylome of the painted lady using whole-genome bisulphite sequencing of experimental cohorts exposed to different environmental cues that potentially can affect the propensity of individuals to allocate resources to migration or reproduction. We combine the methylation data with gene expression profiling of the same samples to uncover the relationship between methylation and gene expression at an unprecedented scale in butterflies. Furthermore, we test if DNA methylation can be a mechanism to transfer environmental signals experienced during larval development to the imaginal stage in female butterflies by splitting larvae into three treatment groups with different larval densities and food availabilities: 1) HDAL (high density, *ad libitum* food), 2) LDAL (low density, *ad libitum* food) and, 3) HDLI (high density, limited food). Since we hypothesize that neurogenetics has a major role in the “decision-making”, we analyse heads (including antennae), where the nervous system of butterflies is concentrated.

## 2. Results

### (a) DNA methylation is concentrated to genic regions

We extracted DNA from the head of 9 adult painted ladies and sequenced their methylome using bisulphite sequencing. Median read depth after deduplication ranged from 9-28X (Table S1). The genome-wide CpG methylation level in *V. cardui* in all samples was low (~ 0.8%). However, a detailed analysis of variation in methylation across genomic regions revealed a considerably higher level in genic regions; both coding sequences (CDS), 3’ untranslated regions (UTRs) and promoters had ~ 10 % methylation (Figure 1). We found that the levels of methylation also varied within gene regions with a considerable dip in methylation around the transcription start site (TSS; Figure 2). Most transposable elements (TEs) had negligible levels of methylation, indicating that methylation is not a major silencing mechanism of TE expression in the painted lady (Figure 1). Overall methylation levels varied between treatments where more stressful conditions resulted in slightly higher methylation levels, especially in promoters, 3’UTRs and CDS (Figure 1). See section (c) for a detailed analysis of differential methylation between treatments.

**Figure 1.**
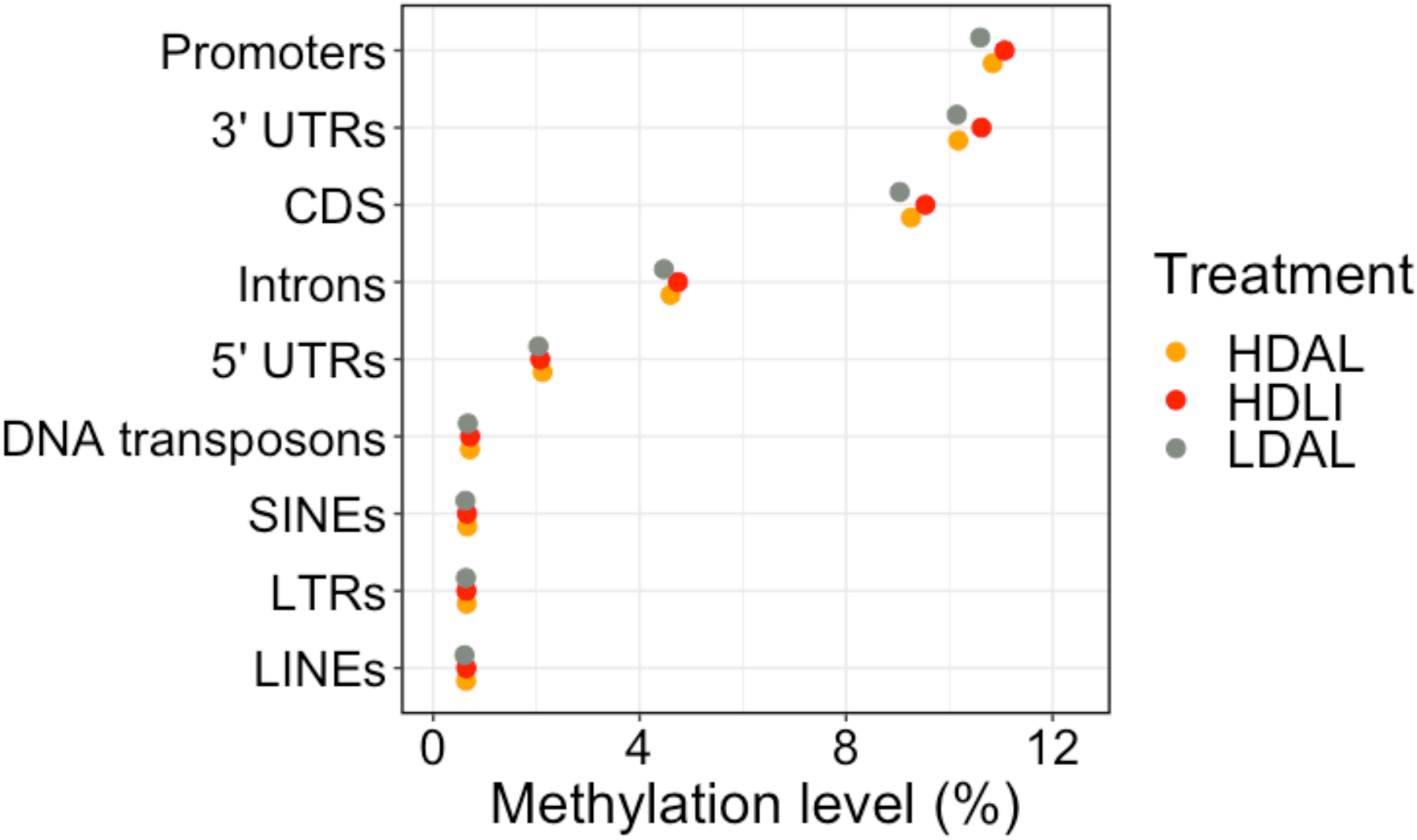
Average methylation levels within different genomic elements for each treatment group.

**Figure 2.**
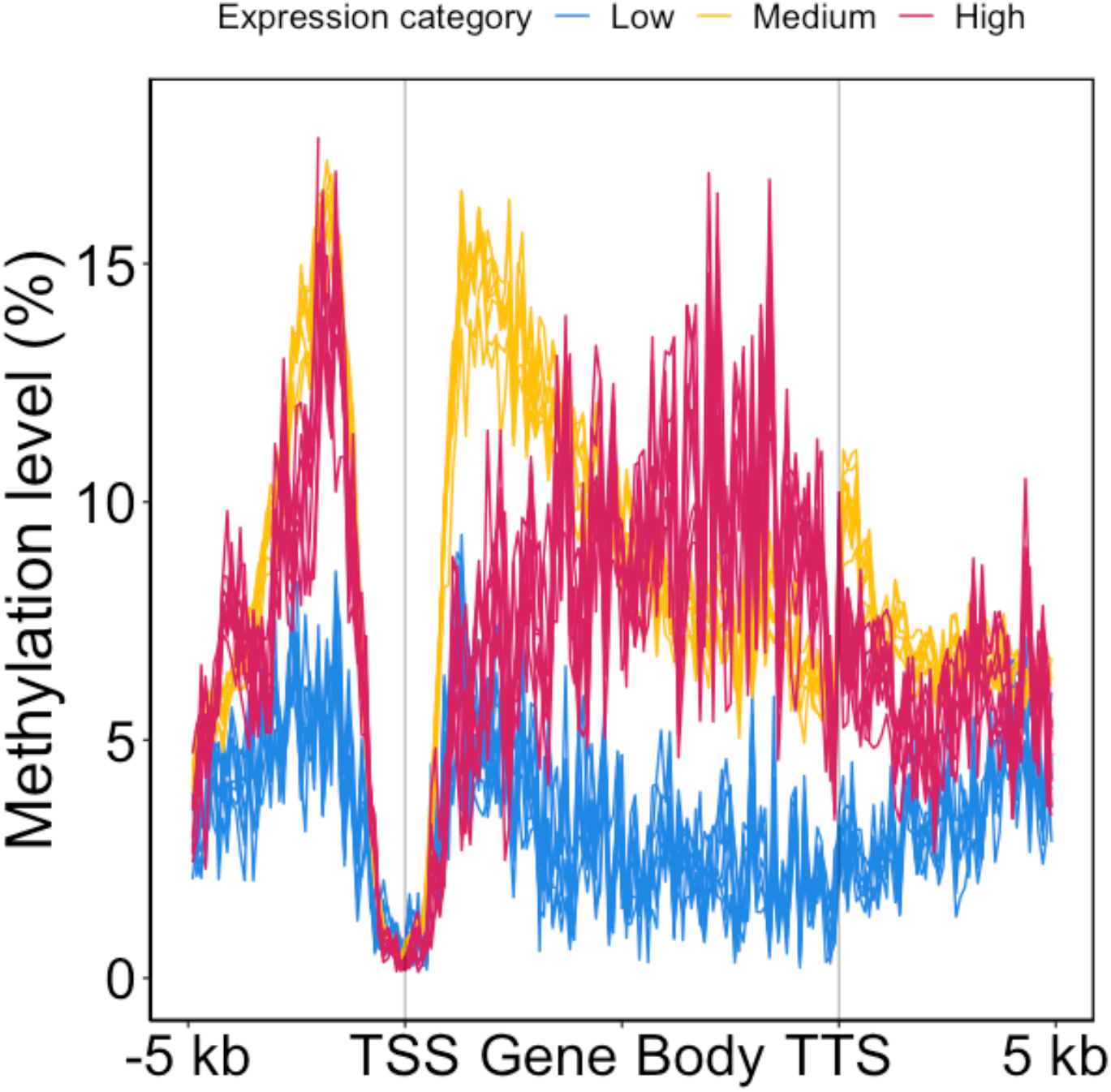
Methylation profiles of all genes with UTR annotation. The three categories represent genes with low (blue; 0-10^th^ percentile), medium (yellow; 10-90^th^ percentile) and high (red; 90-100^th^ percentile) expression levels. TSS and TTS are the inferred transcription start- and termination sites, respectively.

### (b) Methylation levels along the gene body vary with gene expression levels

To assess potential relationships between gene expression and methylation we sequenced the transcriptome of eight of the same samples for which methylation levels were scored and quantified the gene expression levels in each individual. Genes were divided into three rank categories based on expression level: low (0-10^th^ percentile), medium (10-90^th^ percentile) and high (90-100^th^ percentile). All genes, regardless of expression level, were unmethylated close to the TSS (Figure 2). Genes with a higher expression level also had higher methylation levels in the promoters and the gene bodies. The relationship was however quadratic just downstream of the TSS, with intermediate expression levels showing the highest degree of methylation.

### (c) Most differentially methylated regions show higher methylation levels under stress

To identify candidate regions associated with environmental conditions, we performed a genome-wide search for differentially methylated regions (DMRs). Here, two contrasts were used: 1) between the treatments HDAL and LDAL to find regions associated with larval density in *ad libitum* food conditions, and 2) between the treatments HDAL and HDLI, to find regions associated with larval food limitation in high density conditions. Most of the ~ 3,000 DMRs in each contrast showed a higher methylation level in the more stressful condition (Binomial tests *p* < 0.0005; Table S2; Figure 3). We also analysed where in the genome DMRs were localized and found a significant enrichment (Monte-Carlo *p* < 0.003, 1000 replicates) within different gene regions (Figure S1), with 3’ UTRs, CDSs and promoters particularly enriched. This is expected to some extent given the higher average methylation level in these regions which translates to more opportunities for substantial changes in methylation levels.

**Figure 3.**
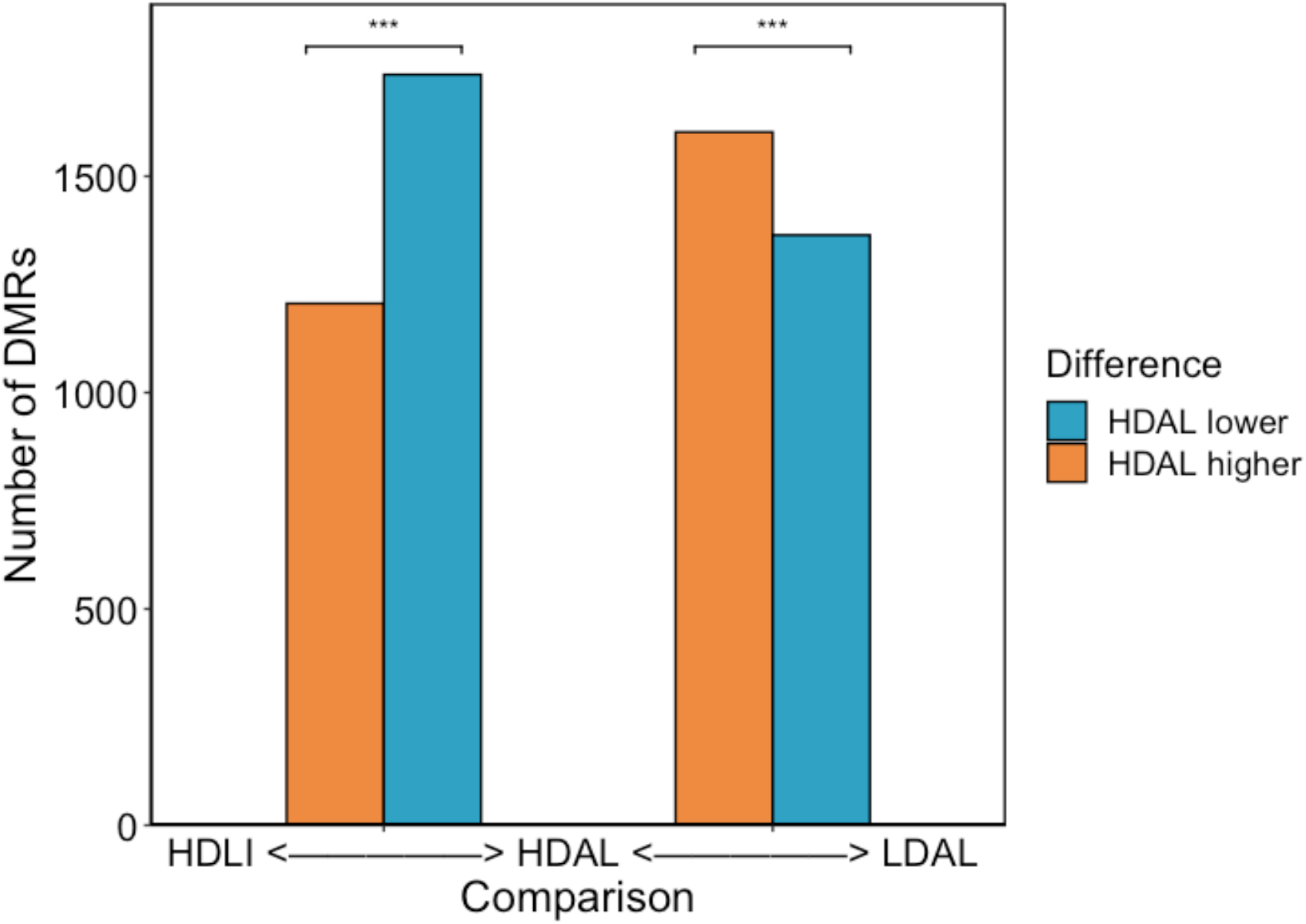
Differentially methylated regions in the two contrasts HDAL-LDAL and HDAL-HDLI, respectively. Treatment groups are sorted based on level of stress, with the most stressful condition to the left (HDLI) and the least stressful (LDAL) to the right. Note that the most stressful condition in each contrast shows a significant (***: *p* < 0.0005) excess of differentially methylated regions (DMRs). HDAL is used as the reference for each contrast.

### (d) Functional analysis of genes enriched in differentially methylated regions between treatments reveals candidate pathways and genes involved in environmental response

We performed a gene ontology enrichment (GO) analysis to investigate whether DMRs overlapping gene features and promoters were associated with genes involved in certain biological processes. For DMRs overlapping CDSs in both comparisons we found enrichment of genes involved in peptide transport (Table S3-4). In addition, for the HDAL-LDAL contrast, we also found enrichment of genes involved in several processes related to translation (Table S4).

Because of the considerably higher methylation level, and consequently a higher power to detect differences, there is a higher risk of detecting false positive DMRs and DMRs without functional consequences in gene regions. Therefore, we focused on genes with multiple DMRs, which are more likely to represent biologically relevant categories, i.e., there is a low probability that many DMRs appear in the same gene by chance. Under the conservative assumption that all identified DMRs overlap CDSs, the expected frequency is ~ 0.2 DMRs per gene. We selected genes with more than four DMRs overlapping the coding sequence in each comparison and in this set we found several genes with known expression in head, antennae, and the central nervous system in *Drosophila* (Table S5-6). In the HDAL-HDLI comparison, we found two candidate genes involved in flight behaviour: *rigor mortis*, which regulates mRNA splicing, and *mettl3*, an enzyme with the ability to methylate adenosines in mRNA (Table S6).

### (e) Differentially expressed genes are not enriched in differential methylation

Differential expression analyses were performed to investigate if the larval treatments resulted in transcriptional profile shifts and whether they were associated with DMRs. In both contrasts, 35 genes were differentially expressed (DE). Eight of those genes were DE in both contrasts. Subsequent GO analysis based on DE genes revealed no significant enrichment of any specific gene categories for the HDAL-LDAL comparison. In the HDAL-HDLI contrast, several biological processes related to translation and nonsense-mediated decay of mRNA were significantly enriched (Table S7). Translational regulation was also found in the GO analysis of DMRs but in the HDAL-LDAL contrast, perhaps indicating some functional concordance with DMRs. The differentially methylated genes *rigor mortis* and *mettle3* were not significantly DE. To test whether DE genes were enriched in DMRs, which is expected if DMRs in *cis* (locally) are the main regulatory cause of DE genes or correlated with a change in expression, we investigated if there was a significant overlap between DE and gene body DMRs. Note that differential methylation is an epigenetic mark that can persist over ontogenetic stages while DE is more of an ephemeral signal (see *Discussion*) which adds some uncertainty to the inference [26,42]. Furthermore, our data do not allow to investigate a potential influence of differential methylation in *trans* (at a location far away from the gene or at another chromosome). The few overlapping regions we identified did not significantly (Fisher’s exact test; *p* > 0.05) deviate from the random expectation (Table 1). In addition, we found no DMRs in promoter regions of DE genes. Thus, we conclude that DE does not always give rise to DMRs in gene bodies of DE genes in *V. cardui* and other regulatory mechanisms are likely to be involved in addition to differential methylation.

**Table 1:** Overlap between DMRs and DE genes. Parentheses in the first column show the 95 % confidence interval in a Fisher’s exact test.

| Comparison | Odds ratio | #DE genes with gene body DMRs | #DE genes | #Genes with DMRs |
| --- | --- | --- | --- | --- |
| HDAL-LDAL | 0.54 (0.11 - 1.71) | 3 | 35 | 2,091 |
| HDAL-HDLI | 0.74 (0.19 – 2.09) | 4 | 35 | 2,020 |

## 3. Discussion

In this study we characterised the DNA methylome of a nearly cosmopolitan migratory butterfly species and tested whether methylation could be a mechanism to store distinct environmental signals during larval development.

### (a) Diversification of methylation within Lepidoptera revealed by novel relationship with expression

Intriguingly, our results showed that the association between methylation and gene expression in *V. cardui* is different from what has previously been observed in *B. mori* [32], indicating that DNA methylation patterns has diversified within Lepidoptera. Methylation across the gene body has been shown to be positively associated with gene expression in *B. mori*, while we observed a more complex pattern in *V. cardui*, with a negative quadratic relationship in the 5’ end of genes and a positive linear relationship at the 3’ end. Our results support that the gene body methylation level in *V. cardui* can be determined by the occupancy level of RNA polymerase, as has been hypothesized to be the case in humans [31]. According to this model, the opportunity of methylation should be greatest when chromatin is open and RNA polymerase occupancy is low, which likely reflects the situation in genes with intermediate expression. For highly expressed genes, the chromatin is probably also highly accessible, but RNA polymerase occupancy levels are high enough to prevent DNA methyltransferase activity. Evidence from several studies indicate that RNA polymerase occupancy is especially high at 5’ end of expressed genes [43], which could explain the observed shifting patterns of methylation levels between 5’ and 3’ ends of the gene body for different levels of gene expression in *V. cardui*. Several hypotheses could explain why the previous studies on *B. mori* may have failed to detect the complex pattern here described for *V. cardui:* 1) There is a difference between these lineages in the mechanism regulating methylation. 2) The mechanism varies according to the tissues analysed. Only the silk gland of instar V larvae was analysed in *B. mori*, while heads of adult butterflies were used here. 3) The pattern varies according to the different gene sets used. Here we restricted the analysis to the 3,685 genes with annotated 5’ and 3’ UTRs to increase accuracy in the inference of TSS and TTS. As previously mentioned, the function of gene body methylation is still debated [29,30]. Targeted partial demethylation at three gene body loci in a *B. mori* embryonic cell line lead to slightly increased levels of gene expression (1.2-1.7 fold) compared to controls [36]. While only at a few loci, these results challenges earlier *B. mori* results [32,33], but in line with the quadratic relationship between DNA methylation and gene expression we observed here among genes. It is also possible that the relationship among genes does not reflect the relationship between treatments at a single gene. More data is needed before any decisive conclusions can be drawn on the function(s) of gene body methylation in Lepidoptera. We also observed more variation in methylation levels in the promoters of *V. cardui* genes compared to *B. mori* genes which appear to predominantly be unmethylated [32]. Consequently, our results show that general promoter methylation levels differ significantly between species/tissues in Lepidoptera. The functional importance of this difference remains to be determined.

### (b) Stress during development shifts the methylation profile in adults but evidence for a relationship with differential expression remains elusive

Our results show that environmental signals during larval development can be stored as methylation patterns in the adult. Further confidence for this conclusion comes from the fact that several genes showing multiple DMRs frequently have been shown to be expressed in homologous structures of the head in *Drosophila melanogaster* [44]. Both treatments that induced stress, especially the HDLI treatment, resulted in an excess of methylation in the identified DMRs. This could be an effect of a prolonged development time induced by stress [45], or a more direct effect on e.g., gene expression causing correlated changes in methylation. The identified DMRs could represent regions that either affect regulation at the time point of sampling, or at any previous time point during development. In contrast, the level of gene expression is likely a much more ephemeral signal [26,42]. Thus, an absence of overlap enrichment between DE and DMRs does not mean that differential methylation has not played a regulatory role. The relationship between gene expression and methylation could alternatively be dominated by *trans* changes or changes at distal regulatory sites. Effects of methylation at *cis*-regulatory elements is transcription-factor dependent in humans [46], and largely unexplored in arthropods [37]. Both the DMRs and DE analyses revealed a concordance with enrichment of genes associated with translational regulation, albeit in two different contrasts that include the HDAL treatment group (see below for a discussion on experimental design). Interestingly, methylated genes in *B. mori*, honey bee (*Apis mellifera*) and fire ant (*Solenopsis* invicta) are also enriched in ontologies related to translation [32,39]. Whether or not methylation has a causal role in determining gene expression levels, the concordance on the level of GO between DMRs and DEs, indicates that they are co-regulated to some extent.

### (c) Differential methylation in an RNA methyltransferase gene: *mettl3*

An interesting candidate gene for further study focusing on the interplay between expression and DNA methylation is *mettl3*, in which we found 4 DMRs overlapping CDS in the HDAL-HDLI contrast. A recent study [47] using human cell cultures showed that *mettl3*-mediated RNA methyladenosine formation leads to demethylation of DNA in neighbouring genomic loci of methylated transcripts, which affects chromatin accessibility and gene expression. If a similar mechanism acts in butterfly cells, it is possible that *mettle3* autoregulates by methylating its own transcript, which in turn leads to increased DNA demethylation at the *mettle3* locus. However, *mettl3* was not DE in the HDAL-HDLI contrast, indicating that differences in *mettle3* activity caused by food availability probably occurs prior to the adult stage.

### (d) Caveats

Separation of the marginal effects of density and food limitation was not possible in our experimental design. For example, in both the HDAL-LDAL and HDAL-HDLI comparisons we observed that peptide transport function was enriched for genes with CDS DMRs. This could be due to similar molecular systems being involved in responses to density (in *ad libitum* food conditions) as well food limitation (in high density conditions), but to confidently assess this we would need to investigate marginal effects of each treatment which requires a 2 x 2 factorial design. We also cannot exclude the effects of the same HDAL samples being reused in both comparisons. For example, 8 of 35 genes were DE in both contrasts as well as 513 of ~ 3,000 DMRs (when requiring > 80% reciprocal overlap). These subsets could either represent true biological signals or correlated false positives in both comparisons. However, it’s reassuring for our conclusions that this set represents a minority of DE genes and DMRs.

### (e) Outlook

Whether the methylation and expression differences between treatment groups we observe here influence migratory behaviour remains to be proven. However, we have shown, as a first step, that high density and food limitation induce molecular programs in the heads of larvae that echoes throughout development. We did uncover two interesting candidate genes, both involved in flight behaviour in *Drosophila melanogaster*. Future studies should investigate their expression profiles across ontogenetic stages and assess their potential functions associated with migratory behaviour using RNAi or CRISPR-Cas9 methodologies.

## 4. Conclusion

Here we provide a detailed characterisation of genome-wide methylation profiles in the painted lady butterfly, a key species for insect migration research. Besides giving a broad sense of how methylation is distributed in a butterfly, we also show that gene-body methylation is variably associated with gene expression in Lepidoptera. Most importantly, we contrast multiple treatment groups and identify differentially methylated regions that provide a first glimpse into how environmental cues can alter the epigenetic landscape and be transmitted from larval to adult stages. The identified candidate regions will provide important starting points for investigating potential mechanistic links between epigenetic modifications and investment in migration or reproduction.

## 5. Materials and methods

### Experimental setup

Mated female painted lady (*Vanessa cardui*) butterflies were collected in Catalonia, Spain, in spring 2020 and kept in individual cages at 25°C, 18:6 hours light:dark regime, with access to a host plant (*Malva sylvestris*) and 10% sugar water. F_1_ offspring were raised on an *ad lib* (AL) *M. sylvestris*, and individually marked as adults and released in a monitored common cage with other F_1_ adults. F_1_ females were separated after first mating and put in individual flasks with *M. sylvestris* for egg-laying. F_2_ eggs from each mated female were split into three different treatment groups representing larval environmental conditions: LDAL (low density, AL food), HDAL (high density, AL food), and HDLI (high density, limited food). LD represented larvae kept individually, and HD represent larvae raised in groups of 10. In the LI treatment, food was only replaced every other day, leading to a 1:1 day food:starvation regime. Female F_2_ adults were flash frozen in liquid nitrogen immediately after eclosure as adults and stored at -80°C until nucleic acid-extraction. The heads (including antennae) of three pairs of siblings in the three treatment groups (Table S1) were homogenized on ice and separated in aliquots for RNA and DNA extraction, respectively.

### DNA extraction and bisulphite sequencing

DNA was extracted using a standard phenol-chloroform extraction protocol. Library preparation and whole-genome bisulphite sequencing were performed by the SciLife SNP&SEQ Technology Platform in Uppsala, Sweden. Sequencing libraries were prepared from 4-200 ng of DNA according to the SPLAT method [48]. Prior to bisulphite conversion, 0.5-1% of lambda phage was spiked-in. The bisulphite conversion error rate was low and varied from 0.2-0.3 % between samples. Paired-end sequencing (150 bp read length) was performed using the NovaSeq 6000 technology and v1.5 sequencing chemistry in a single S4 flow-cell.

### RNA extraction and sequencing

The RNeasy Mini Kit (Qiagen) was used to extract RNA following the general instructions in the kit manual. Library preparations (Illumina TruSeq Stranded mRNA polyA selection kit) and RNA sequencing were performed by the National Genomics Infrastructure (NGI) in Stockholm. Libraries were sequenced on two lanes on one S4 flow-cell on the NovaSeq S6000 with 150 bp paired-end reads (in multiplex with additional samples used for another study, data not shown).

### Processing of bisulphite sequence reads and methylation calling

All processing steps from adapter filtering to read mapping, and methylation calling was performed using the reproducible Nextflow v21.04.1 nf-core Methylseq v1.5 pipeline [49]. Reads were trimmed using Trim Galore! v0.6.4_dev, employing Cutadapt v.2.9 [50]. Further clipping of reads was performed using the Epignom profile (8 bp from both 5’ and 3’ ends of both reads in a pair). Trimmed reads were aligned using Bismark v0.22.3 [51] to a previously available *Vanessa cardui* reference genome [52]. Bismark was also used for methylation calling.

### Gene expression quantification and differential expression analysis

All processing steps from adapter filtering to read mapping and transcript quantification was performed using the reproducible Nextflow v21.10.6 nf-core rnaseq v3.8.1 pipeline [49]. Reads were trimmed using Trim Galore! v0.6.7, employing Cutadapt v.3.4 [50]. Reads were mapped to the reference using STAR v2.7.10a [53] and a previously available gene annotation consisting of 13,161 genes [15]. Read quantification was performed using salmon v1.5.2 [54], and we used transcripts per million reads (TPM) values as a measure of gene expression level. Differential expression analyses was performed using deseq2 v1.28.0 in R v.4.2.1 taking into account sibling relationship using the design: ~Family+Treatment [55]. A false discovery rate of 0.1 was used to correct for multiple testing.

### Methylation level

When using bulk samples of DNA, we need to measure methylation as a proportion, since chromatids differ in methylation status dependent on cell type and methyltransferase dynamics. To be precise, it is the combined level of 5’-CpG-3’ methylation and hydroxymethylation that is measured since bisulphite sequencing cannot distinguish these marks [56]. Here we measured methylation level by summarizing the number of methylated *x_mCpG_* reads mapping to both strands of a reference CpG dinucleotide and dividing by the total number of reads *x_uCpG_* + *x_mCpG_*. To measure the methylation level covering *n* CpG dinucleotides, we used the average proportion across the individual dinucleotides (*i*),

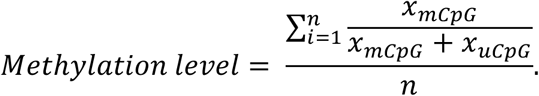

### Gene profile

We selected genes having at least one transcript with 5’- and one with a 3’UTR, such that an average TSS and TTS could be defined with more confidence. This retained 3,685 out of 13,161 annotated genes. Regions up- and downstream of each gene were split into 100 bp non-overlapping segments. Gene bodies were also split into 100 bp segments, then normalised jointly in 99 windows based on rank along the gene profile.

### Differential methylation analysis

We identified DMRs using BSmooth [57] and defined them as regions at the top two percentile of genome-wide absolute methylation differences, keeping only CpG sites where at least two samples each, for both groups had a coverage >= 2. We further required DMRs to contain at least three CpGs with a mean methylation difference >= 10 percentage points. We used a resampling method to shuffle the DMRs *n* =1,000 times across the genome to obtain an expected amount of overlap with annotated features. We calculated a two-tailed empirical *p*-value as 2*r*/*n* for *r/n* <= 0.5 and 2(1- *r/n*) for *r/n* > 0.5, where *r* is the number of replicates with an overlap greater than or equal to the overlap for the observed data [58]. Enrichment was defined as the following odds ratio:

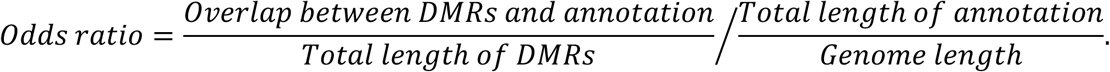

### Gene ontology enrichment analysis

All gene ontology enrichment analyses were performed using the “weight01” algorithm in topGO [59]. We used the Benjamini-Hochberg method to correct for multiple-testing to a family wise error rate (FWER) of 0.1.

### Statistical analysis

Statistical analyses except permutations were performed in R v.4.1.3 and/or v.4.2.1 [60].

## Data access

Bisulphite and RNA-seq data are available at the European Nucleotide Archive under study id ERP142326. Scripts are available on Github in the following repository: https://github.com/JesperBoman/DNA-methylation-of-the-Painted-Lady.

## Contributions

JB – wrote and revised the original manuscript, experimental design, data analysis

NB – writing, experimental design, funding, lab work

LH – writing, lab work, experimental design

YZ – writing, lab work, experimental design

GT – writing, funding, data acquisition

RV – writing, funding

## Competing interest statement

The authors declare no competing interests

## Acknowledgements

This project was financially supported by a FORMAS research grant (2019-00670 to N.B.). Bisulphite sequencing was performed by the SNP&SEQ Technology Platform in Uppsala. The facility is part of the National Genomics Infrastructure (NGI) Sweden and Science for Life Laboratory. The SNP&SEQ Platform is also supported by the Swedish Research Council and the Knut and Alice Wallenberg Foundation. RNA-sequencing was performed by the National Genomics Infrastructure in Genomics Production, Stockholm, funded by the Science for Life Laboratory, the Knut and Alice Wallenberg Foundation and the Swedish Research Council. We thank SNIC/Uppsala Multidisciplinary Center for Advanced Computational Science for access to the UPPMAX computational infrastructure and acknowledge the National Bioinformatics Infrastructure Sweden (NBIS) for granting WABI (Wallenberg Advanced Bioinformatics Infrastructure) support to the research group. R.V. was supported by the Spanish government through grant PID2019-107078GB-I00/ MCIN/AEI/ 10.13039/501100011033. G.T. was supported by the project PID2020-117739GA-I00 MCIN / AEI / 10.13039/501100011033. All authors were supported by the grant LINKA20399 from the CSIC iLINK program. We thank Elenia Parkes for contributing to RNA extractions, under supervision of Lars Höök. The authors also thank Erik Gudmunds and Karin Näsvall for insightful discussions about the results.

## Notes

### Competing Interest Statement

The authors have declared no competing interest.

