## Supplementary Information for "Environmental stress during larval development induces head methylome profile shifts in the migratory painted lady butterfly (*Vanessa cardui*)"

**Table S1**: Sequencing and sample information. Bisulfite- (BS) and RNA-seq reads are provided in million reads.

| **Sample** | **Family** | **Treatment** | **BS-seq reads** | **Median coverage mapped BS-seq reads** | **RNA-seq reads** |
| --- | --- | --- | --- | --- | --- |
| LDAL-X1 | X1 | LDAL | 171.4 | 26.0 | 70.7 |
| LDAL-X3 | X3 | LDAL | 86.5 | 12.0 | 62.1 |
| LDAL-X18 | X18 | LDAL | 51.7 | 9.0 | 91.6 |
| HDAL-X1 | X1 | HDAL | 125.0 | 19.0 | 88.6 |
| HDAL-X3 | X3 | HDAL | 152.2 | 28.0 | 86.8 |
| HDAL-X18 | X18 | HDAL | 72.8 | 11.0 | 81.1 |
| HDLI-X1 | X1 | HDLI | 163.4 | 28.0 | NA * |
| HDLI-X3 | X3 | HDLI | 71.7 | 12.0 | 60.4 |
| HDLI-X18 | X18 | HDLI | 81.3 | 13.0 | 62.3 |

***** RNA extraction of sufficient quality failed for this sample

**Figure S1:** Odds ratio of overlap between DMRs and annotated features. Significance was determined using 1,000 Monte Carlo replicates with a family wise error rate (FWER) of 0.1.

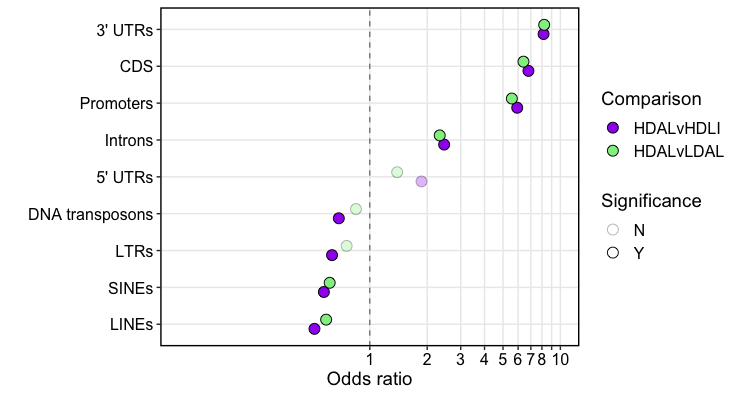

**Table S2**. Characteristics of methylation contrasts between treatment groups. Both nutritional (food limitation) and social stress (higher larval density) generally led to an increase in methylation. Parentheses in the third column show the 95 % confidence interval in a binomial test.

| **Comparison** | **Number of DMRs** | **Proportion of DMRs with higher methylation in HDAL** |
| --- | --- | --- |
| HDAL-LDAL | 2966 | 0.54 *** (0.52-0.56) |
| HDAL-HDLI | 2941 | 0.41*** (0.39-0.42) |

*** *p <* 0.0005

**Table S3**: GO analysis of genes with DMRs overlapping CDS in the HDAL-LDAL comparison for the biological process category. *P-*values were adjusted for multiple testing using the Benjamini-Hochberg method to an FWER of 0.1.

| **GO ID** | **Term** | **Annotated** | **Significant DMR** | **Expected** | **Adjusted *p*** |
| --- | --- | --- | --- | --- | --- |
| GO:0006614 | SRP-dependent cotranslational protein targeting to membrane | 56 | 22 | 6.82 | 0.001639555 |
| GO:0015833 | Peptide transport | 1107 | 189 | 134.75 | 0.001639555 |
| GO:0006413 | Translational initiation | 138 | 40 | 16.8 | 0.007603733 |
| GO:0002181 | Cytoplasmic translation | 113 | 36 | 13.76 | 0.008554200 |
| GO:0019886 | Antigen processing and presentation of exogenous peptide antigen via MHC class II | 57 | 19 | 6.94 | 0.068433600 |
| GO:0000184 | Nuclear-transcribed mRNA catabolic process, nonsense-mediated decay | 86 | 25 | 10.47 | 0.076037333 |

**Table S4**: GO analysis of genes with DMRs overlapping CDS in the HDAL-HDLI comparison for the biological process category. *P-*values were adjusted for multiple testing using the Benjamini-Hochberg method to an FWER of 0.1.

| **GO ID** | **Term** | **Annotated** | **Significant DMR** | **Expected** | **Adjusted *p*** |
| --- | --- | --- | --- | --- | --- |
| GO:0015833 | Peptide transport | 1107 | 178 | 141.69 | 0.057028 |

**Table S5:** Genes with more than four DMRs in the CDS in the HDAL-LDAL comparison.

Functional information from gene annotation (ref), GeneCards and/or Flybase.

| **Number of DMRs** | **Gene** | **Gene function** |
| --- | --- | --- |
| 6 | CCZ1 | Enables guanyl-nucleotide exchange factor activity. Predicted to be involved in vesicle-mediated transport. Located in intracellular membrane-bounded organelle. |
| 5 | WARS1 | Tryptophanyl-TRNA synthetase |
| 5 | ? | Zinc finger |
| 5 | SDS3 | Component of the RPD3C(L) histone deacetylase complex (HDAC) responsible for the deacetylation of lysine residues on the N-terminal part of the core histones (H2A, H2B, H3 and H4). |
| 5 | ? | C2H2 zinc finger |
| 5 | SMARCA5 | The protein encoded by this gene is a member of the SWI/SNF family of proteins. Members of this family have helicase and ATPase activities and are thought to regulate transcription of certain genes by altering the chromatin structure around those genes. |
| 4 | MRPL37 | Predicted to be a structural constituent of ribosome. Predicted to be involved in mitochondrial translation. Predicted to be part of mitochondrial large ribosomal subunit. Predicted to be active in mitochondrion. Is expressed in several structures, including anterior endoderm; anterior endoderm anlage; dorsal pharyngeal muscle cell; extended germ band embryo; and gut section. |
| 4 | ING3 | Predicted to enable methylated histone binding activity. Involved in histone acetylation and histone exchange. Located in nucleus. Part of NuA4 histone acetyltransferase complex. |
| 4 | NUP154 | Enables chromatin binding activity. Involved in several processes, including male gamete generation; oogenesis; and positive regulation of protein import into nucleus. Located in cytoplasm; nuclear envelope; and nuclear periphery. Part of nuclear pore. Is expressed in several structures, including embryonic/larval circulatory system; embryonic/larval gut; gastrula embryo; gonad; and presumptive embryonic/larval central nervous system. |
| 4 | Rabankyrin-5 | Predicted to enable metal ion binding activity. Involved in embryonic morphogenesis; endocytosis; and epithelial cell morphogenesis. Located in vacuole. Is expressed in adult head; head mesoderm; organism; trunk mesoderm; and ventral ectoderm. |
| 4 | SCO1/Scox | Synthesis of cytochrome c oxidase (Scox) encodes a protein involved in cytochrome complex assembly and regulation of ATP biosynthesis. |
| 4 | GHITM | Involved in inner mitochondrial membrane organization and negative regulation of release of cytochrome c from mitochondria. Located in mitochondrion. Is integral component of mitochondrial inner membrane. |
| 4 | Gawky/TNRC6C | Required for gene silencing mediated by micro-RNAs (miRNAs). Silences both polyadenylated and deadenylated mRNAs. Required for miRNA-mediated translational repression and mRNA decay. Not required for miRNA target recognition. Necessary to initiate but not to maintain silencing. Promotes mRNA deadenylation through the recruitment of the CCR4-NOT and PAN complexes and promotes decapping by the DCP1-DCP2 complex. Dissociates from silenced mRNAs after deadenylation. |
| 4 | UBE3A | Ubiquitin protein ligase E3A (Ube3a) encodes the founding member of the HECT-type ubiquitin E3 ligase family of enzymes. It is involved in the final step of conjugation of ubiquitin to its target substrates. It regulates protein degradation by targeting modified proteins to the proteasome or by regulating the proteasome activity through ubiquitination of its subunits, which in turn affects many aspects of neuronal function, such as synaptic plasticity, long-term memory or dendritic development. |
| 4 | SMYD5 | Predicted to enable S-adenosylmethionine-dependent methyltransferase activity. Predicted to be involved in histone lysine methylation. Predicted to be located in chromatin. Is expressed in anterior midgut primordium; embryonic/larval midgut; embryonic/larval muscle system; organism; and posterior midgut primordium. |
| 4 | IPO5 | Nucleocytoplasmic transport, a signal- and energy-dependent process, takes place through nuclear pore complexes embedded in the nuclear envelope. The import of proteins containing a nuclear localization signal (NLS) requires the NLS import receptor, a heterodimer of importin alpha and beta subunits also known as karyopherins. Importin alpha binds the NLS-containing cargo in the cytoplasm and importin beta docks the complex at the cytoplasmic side of the nuclear pore complex. |
| 4 | NUP54 | A structural constituent of nuclear pore. Involved in NLS-bearing protein import into nucleus. Predicted to be part of nuclear pore central transport channel. Is expressed in several structures, including embryonic central brain neurons; embryonic/larval fat body; extended germ band embryo; germ layer; and gut section. |
| 4 | SNUPN | The nuclear import of the spliceosomal snRNPs U1, U2, U4 and U5, is dependent on the presence of a complex nuclear localization signal. |
| 4 | CSN1a | Predicted to be involved in protein deneddylation. Predicted to be part of COP9 signalosome. |
| 4 | EIFG3 | The protein encoded by this gene is thought to be part of the eIF4F protein complex, which is involved in mRNA cap recognition and transport of mRNAs to the ribosome. |
| 4 | GPR107 | Predicted to enable clathrin heavy chain binding activity. Predicted to be involved in clathrin-dependent endocytosis. Located in Golgi apparatus and nucleoplasm |

**Table S6:** Genes with more than four DMRs in the CDS in the HDAL-HDLI comparison.

Functional information from gene annotation (ref), GeneCards and/or Flybase.

| **Number of DMRs** | **Gene** | **Gene function** |
| --- | --- | --- |
| 7 | ? | Chromatin organization modifier domain |
| 6 | Rigor mortis /GEMIN5 | Involved in flight behavior; molting cycle; and response to ecdysone. This gene encodes a WD repeat protein that is a component of the survival of motor neurons (SMN) complex. The SMN complex plays a critical role in mRNA splicing through the assembly of spliceosomal small nuclear ribonucleoproteins (snRNPs), and may also mediate the assembly and transport of other classes of ribonucleoprotein. |
| 5 | FUCA1 | Alpha-L-fucosidase |
| 4 | Fibrillarin | Predicted to enable RNA binding activity; histone-glutamine methyltransferase activity; and rRNA methyltransferase activity. |
| 4 | METTL3 | Methyltransferase like 3 (Mettl3) encodes an RNA N6-methyladenosine transferase that contributes to oogenesis, lifespan and behaviour, including flight, locomotion and grooming. |
| 4 | PPWD1 | Enables cyclosporin A binding activity and peptidyl-prolyl cis-trans isomerase activity. Involved in protein peptidyl-prolyl isomerization. Located in nuclear body. Part of catalytic step 2 spliceosome. |
| 4 | ELP1 | The protein encoded by this gene is a scaffold protein and a regulator for three different kinases involved in proinflammatory signaling. |
| 4 | DOLK | Predicted to enable dolichol kinase activity. Predicted to be involved in dolichyl monophosphate biosynthetic process. Predicted to be integral component of endoplasmic reticulum membrane. Is expressed in adult head and organism. |
| 4 | BBS1 | Predicted to enable patched binding activity and smoothened binding activity. Involved in cilium assembly. Predicted to be located in cilium. Predicted to be part of BBSome. Predicted to be active in axoneme and centrosome. Is expressed in antenno-maxillary complex; chordotonal neurons; and eo neurons. |
| 4 | ELP2 | Component of the elongator complex which is required for multiple tRNA modifications, including mcm5U (5-methoxycarbonylmethyl uridine), mcm5s2U (5-methoxycarbonylmethyl-2-thiouridine), and ncm5U (5-carbamoylmethyl uridine) (By similarity). |
| 4 | CDH23 | This gene is a member of the cadherin superfamily, whose genes encode calcium dependent cell-cell adhesion glycoproteins. The encoded protein is thought to be involved in stereocilia organization and hair bundle formation |
| 4 | NOC2L | Histone modification by histone acetyltransferases (HAT) and histone deacetylases (HDAC) can control major aspects of transcriptional regulation. NOC2L represents a novel HDAC-independent inhibitor of histone acetyltransferase (INHAT). |
| 4 | SLC44A1 | Enables choline transmembrane transporter activity. Involved in choline transport and transmembrane transport. |
| 4 | TSEN54 | Involved in tRNA processing. Predicted to be part of tRNA-intron endonuclease complex. |
| 4 | CNOT8 | Enables poly(A)-specific ribonuclease activity. Involved in exonucleolytic catabolism of deadenylated mRNA and positive regulation of cell population proliferation |
| 4 | SMARCA5 | The protein encoded by this gene is a member of the SWI/SNF family of proteins. Members of this family have helicase and ATPase activities and are thought to regulate transcription of certain genes by altering the chromatin structure around those genes. |
| 4 | INTS7/deflated | deflated (defl) encodes a component of 'Integrator', a multi-subunit protein complex that binds to the C-terminal domain of RNA polymerase II and is involved in RNA transcript processing and maturation.  Is expressed in embryonic/larval central nervous system; embryonic/larval gut; and organism. |
| 4 | RBM17/Spf45 | Predicted to enable nucleic acid binding activity. Involved in DNA repair; compound eye morphogenesis; and regulation of alternative mRNA splicing, via spliceosome. Part of precatalytic spliceosome. |
| 4 | ARIH2 | The protein encoded by this gene is an E3 ubiquitin-protein ligase that polyubiquitinates some proteins, tagging them for degradation. |
| 4 | LMNB1 | This gene encodes one of the two B-type lamin proteins and is a component of the nuclear lamina. |
| 4 | RASGEF1C | Predicted to enable guanyl-nucleotide exchange factor activity. Predicted to be involved in regulation of catalytic activity and small GTPase mediated signal transduction. |
| 4 | DNAJC11 | Involved in cristae formation. Located in mitochondrial outer membrane and nuclear speck |

**Table S7**: GO analysis of DE genes in the HDAL-HDLI comparison for the biological process category.

| **GO.ID** | **Term** | **Annotated** | **Significant DE** | **Expected** | **Adjusted *p*** |
| --- | --- | --- | --- | --- | --- |
| GO:0006614 | SRP-dependent cotranslational protein targeting to membrane | 55 | 7 | 0.13 | 3.86 * 10^-7^ |
| GO:0000184 | Nuclear-transcribed mRNA catabolic process, nonsense-mediated decay | 86 | 7 | 0.21 | 4.86 * 10^-6^ |
| GO:0002181 | Cytoplasmic translation | 113 | 7 | 0.27 | 2.24 * 10^-5^ |
| GO:0006413 | Translational initiation | 138 | 7 | 0.33 | 6.78 * 10^-5^ |
